# Ultra-High-Throughput LC-MS Method for Targeted Protein Degradation Compound Screening Using the Orbitrap Astral Mass Spectrometer

**DOI:** 10.1101/2025.10.18.683219

**Authors:** Hanfeng Lin, Yen-Yu Yang, Sudipa Maity, Qingyu Sun, Jin Wang

## Abstract

High-throughput proteomics analysis is crucial for targeted protein degradation (TPD) compound screening. Here, we developed a 300 sample per day (SPD) LC-MS method using the Orbitrap Astral mass spectrometer for ultra-high-throughput TPD compound screening. We identified close to 8,000 protein groups from a single cell line with a coefficient of variance (CV) of less than 10%, highlighting the deep proteome coverage and method reproducibility even at 300 SPD. This high degree of precision provides the statistical confidence to detect subtle, yet significant, changes in protein abundance that were previously challenging to quantify in high-throughput workflows. To evaluate the quantitation accuracy of this method, we further mixed the digests from two or three species at different ratios. Our three-proteome mixture results demonstrated highly accurate quantitation for proteins with both small and large fold changes. Moreover, our two-proteome mixture experiment, where 20 to 160 ng of yeast digest was spiked into 200 ng of HeLa digest, showed an R^2^ of 0.999 for the yeast proteome, underscoring the quantitation accuracy of the method. Utilizing this workflow, we studied dose-dependent protein degradation patterns induced by pomalidomide, iberdomide, and mezigdomide. Our results indicate that mezigdomide may possess enhanced efficacy in T cells by degrading additional proteins such as IKZF2, thereby boosting anti-cancer immunity. Together, we developed an ultra-high-throughput LC-MS method with excellent proteome coverage and quantitation accuracy that is highly suitable for chemoproteomics screening of drug libraries.

## Introduction

Mass spectrometry-based proteomics has become an indispensable tool for understanding complex biological systems and for driving drug discovery. In chemoproteomics and targeted-protein degradation (TPD) field, the ability to screen large libraries of compounds against cellular proteomes is critical for identifying and characterizing novel therapeutics. This necessitates the development of high-throughput analytical methods that do not sacrifice depth, precision, or quantitative accuracy ^1–4^. A key challenge has been the reliable detection of subtle protein abundance changes (e.g., 10-20% degradation), as these modest effects are often obscured by analytical variance in conventional high-speed methods.

Over the last five years, data-independent acquisition (DIA) has emerged as a powerful technique for quantitative proteomics with the maturation of data analysis algorithm and software ^5–10^, offering comprehensive peptide detection and reproducible quantification across large sample cohorts. Unlike data-dependent acquisition (DDA), DIA systematically fragments all ions within predefined mass-to-charge (m/z) windows, creating complex but information-rich datasets that can be retrospectively mined ^8,11–14^. However, the adoption of DIA for ultra-high-throughput screening has been historically limited by a trade-off between analytical speed, proteome coverage, and sensitivity. Shortening liquid chromatography (LC) gradients to increase sample throughput often leads to a significant loss in the number of identified and quantified proteins.

The recent introduction of the Orbitrap Astral mass spectrometer marks a significant advancement in addressing these challenges. This innovative platform combines a high-resolution Orbitrap mass analyzer with a novel, high-speed Astral mass analyzer, enabling unprecedented scan speeds (up to 200 Hz) with high resolution, sensitivity and a wide dynamic range ^15,16^. This unique architecture is exceptionally well-suited for DIA, as it allows for the rapid acquisition of high-quality fragment ion spectra even with the narrow chromatographic peaks produced by ultra-short LC gradients ^17,18^. This capability opens the door to achieving both high throughput and high quantitative fidelity simultaneously.

In this study, we leverage the capabilities of the Orbitrap Astral mass spectrometer to develop and validate an ultra-high-throughput LC-MS workflow capable of analyzing 300 samples per day (SPD). We systematically benchmark the quantitative accuracy and precision of the method using standardized multi-proteome mixtures, with a particular focus on its ability to accurately quantify small fold changes against a complex background. We then demonstrate its suitability for deep and reproducible profiling of human cell lines. Finally, we apply this workflow to a real-world TPD study, profiling the dose-dependent effects of several immunomodulatory drugs (IMiDs) to showcase its power for high-throughput compound screening.

## Methods

### Cell Culture and Drug Treatment

Jurkat cell line was purchased from ATCC. Jurkat cells were cultured in RPMI-1640 (Corning, 10-040-CV) supplemented with 10% FBS (Corning, 35-011-CV) in the 37 °C incubator with 5% CO_2_. The day before treatment, 100 μL Jurkat cells were plated in a 96-well U-bottom plate with a density of 2 million/mL. On the following day, a serial dilution of degrader compounds in DMSO or equal volume of DMSO were added into each well, making the final compound concentrations as 8, 40, 200, 1000 nM. Two biological replicates were prepared for each compound concentration and DMSO-treated controls.

### TPD Global Proteomics Sample Preparation

After 24 hours of degrader treatment, cells were centrifuged down, and the media was aspirated, followed by a 1x PBS wash. Cell pellets were lysed in 50 μL of 200 mM HEPES, 0.25% SDS at pH 8.5, with 1x Halt protease inhibitor cocktail (Thermo Scientific, 78440) and 10U/well benzonase (Sigma, E1014), shaking at 800 rpm at 37°C for 30 min. Reduction was performed by adding 5 μL of 50 mM DTT and shaking at 37°C for 15 min, followed by alkylation with 14 μL of 100 mM iodoacetamide at room temperature in the dark for 30 min. To each well, 100 μg of SP3 beads (a 1:1 mixture of E3 and E7 carboxylic magnetic beads from Cytiva) were added. Protein binding was initiated by adding 75 μL of 100% ethanol to each well and shaking at 800 rpm for 10 min at room temperature. The beads were washed twice on a magnetic rack with 200 μL of 80% ethanol. After the final wash, beads were briefly dried, and proteins were digested in 30 μL of 12.5 ng/μL trypsin/LysC (Promega, V5073) in 200 mM HEPES, shaking at 1,000 rpm at 37°C for 18 hours with the plate sealed.

The digested peptides were acidified with 1 μL/well of formic acid (FA) and further desalted and cleaned up with StageTips (CDS Analytical, 6091). Tips were moistened by adding 30 μL of 80% ACN/0.1% FA, centrifuging the assembly at 800 rpm for 1 min, and removing the waste liquid. Air was plunged through to remove excess liquid from the tips. The tips were then equilibrated by adding 30 μL of 0.1% FA and repeating the centrifugation and plunging steps. The acidified digests were loaded onto the StageTips, followed by centrifugation and plunging. The tips were washed twice with 30 μL of 0.1% FA and peptides were eluted with 100 μL of 80% ACN/0.1% FA into the 96-well PCR collection plate. The eluate plate was dried in a speed vac, and peptides were resuspended in 30 μL of 5% ACN/0.1% FA. Peptide concentration was measured with a Fluorescent Quantitative Peptide Assay (Pierce, 23290) and normalized across the plate before LC-MS/MS injection.

### Generation of Proteome Mixtures

Human HeLa cell digest (Thermo Scientific, 88328), E. coli digest (Waters Corporation, 186003196), and yeast digest (Thermo Scientific, A47951) standards were dissolved in 5% ACN/ 0.1% FA as 10 ng/μL stock solutions. For three proteome mix samples, various volumes of digests were combined to make the final sample containing 2 μg total proteomes with the specified percentage of components. For two-proteome mix samples, 20, 40, 60, 80, 100, and 160 ng of yeast digest were spiked into a 200 ng HeLa digest background. Samples were Speed-Vac dried and reconstituted in 10 μL 5% ACN/ 0.1% FA immediately before loading for LC-MS/MS analysis.

### Liquid Chromatography and Mass Spectrometry (LC-MS) Analysis

A total of 50 or 200 ng of the reconstituted peptides were loaded onto a PepMap Neo Trap Cartridge (Thermo Scientific, 174500) using a Vanquish Neo UHPLC system (Thermo Scientific) and separated by a 15-cm EASY-Spray PepMap column (Thermo scientific, ES906) for 60 or 180 SPD, or a 50-cm EASY-Spray PepMap Neo HPLC column (Thermo Scientific, ES75500PN) for 24 SPD analysis. For 300 SPD analysis, the peptides were resolved by an Aurora Rapid® 5×75 C18 UHPLC column or Aurora Rapid® 8×75 XT C18 UHPLC column (IonOpticks). Flow rate and gradients were optimized for individual column and throughput, and the parameters are detailed in **Supplementary Table 1**.

For DIA analysis, MS1 spectra were collected on the Orbitrap mass analyzer with a resolution of 240,000 and a scan range of 380-980 m/z and 580-780 m/z for 24, 60, 180 SPD and 300 SPD, respectively. The AGC was set as 500%, and the maximum injection time was set to 5 ms. Precursor ions were fragmented through HCD with a normalized collision energy (NCE) of 25%. MS2 spectra were acquired on the Astral mass analyzer with the parameters of each of the throughput listed in **Supplementary Table 1**.

### Data Processing and Statistical Analysis

Data was processed by Spectronaut software (Biognosys, v19) using the directDIA approach. The spectra were searched against human, yeast or E. coli proteome downloaded from Uniprot. Enzyme was set as trypsin that cleaves the C-terminal of lysine or arginine except for those immediately after proline. Carbamidomethylating on cysteine was set as fixed modification; acetylation on protein N-terminal and oxidation on methionine was set as variable modification. The FDR control was set as 1% for PSM, peptide and protein group. Normalization was performed by using peptides in human fasta files for all experiments. The search results were exported into tabular formats and imported to Python for downstream data analysis and visualization.

### Data and code availability

All the raw data from Orbitrap Astral has been deposited to MassIVE database under accession: MSV000099486. (PW: g1Yz1PJr6Dd8xJZt)

All the codes for Astral DIA data analysis have been deposited at https://github.com/Hanfeng-Lin/Astral_DIA_analysis.

## Results

### Benchmarking the Orbitrap Astral with Various Throughputs

To establish the performance of the Orbitrap Astral mass spectrometer for our high-throughput proteomics workflow (**Figure 1A**), we first evaluated its quantitative accuracy with both long and short gradients. We analyzed a standard three-proteome mixture (Human, Yeast, *E. coli*, **Figure 1B**) at different throughputs, ranging from 24 to 300 samples per day (SPD) (**Table S1**). The results demonstrated that the instrument is highly suitable for both ultra-high-throughput analysis and longer gradients intended for in-depth proteome coverage. Across all tested gradients, the measured protein ratios for each species remained highly accurate and precise, aligning closely with the expected values (**Figure 1C**). This initial benchmark confirms the instrument’s robust quantitative performance, which is essential for reliable compound screening.

**Figure 1.**
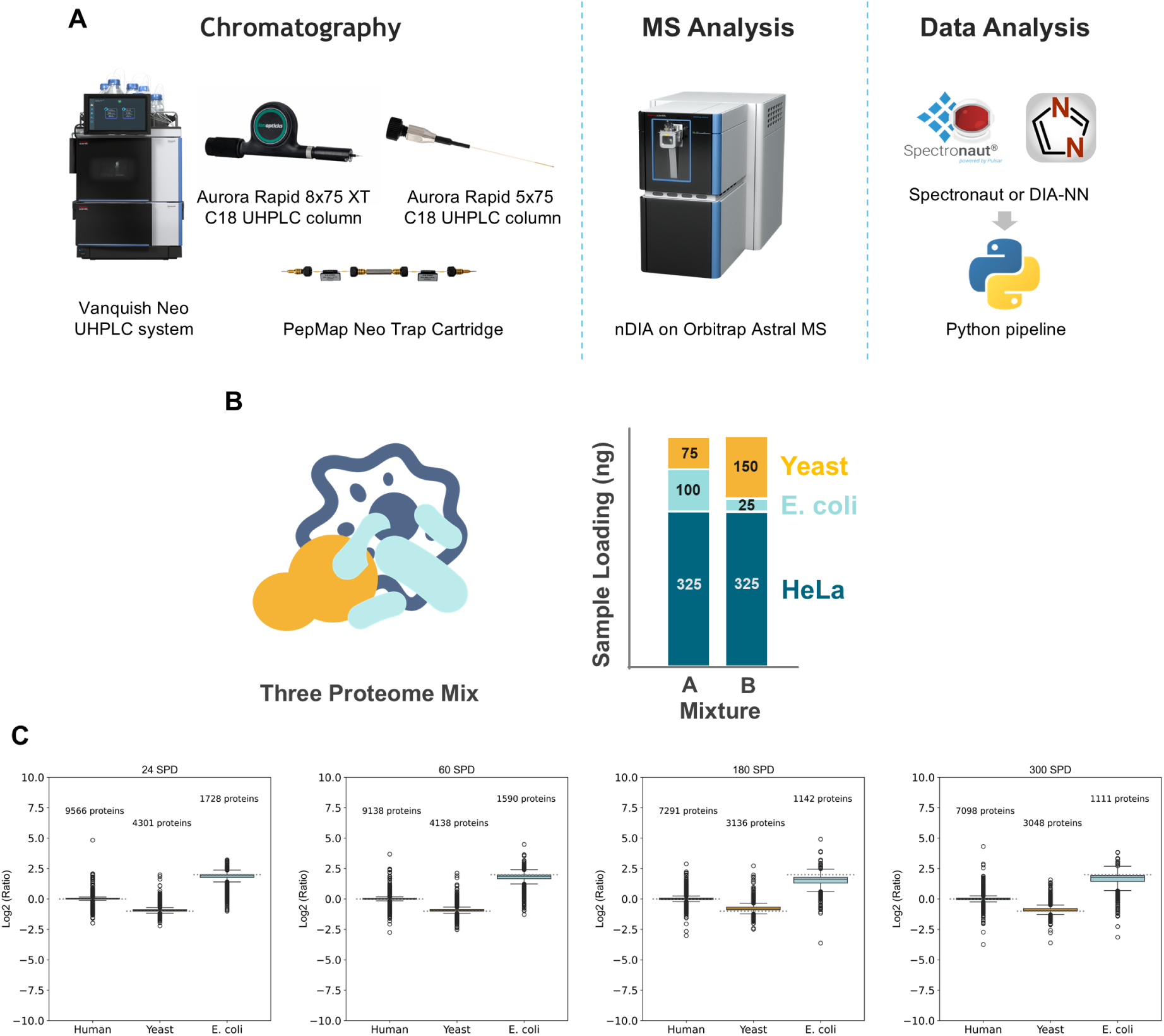
Benchmarking Quantitative Accuracy Across Various Throughputs. (A) Schematic overview of the liquid chromatography-mass spectrometry (LC-MS) workflow. (B) Composition of the standard three-proteome mixture used to assess quantitative performance, with digests from human (HeLa), yeast, and *E. coli* mixed at defined ratios. (C) Box plots showing the measured log_2_Ratio of proteins from each proteome. The analysis was performed at throughputs of 24, 60, 180, and 300 SPD. The dashed lines indicate the theoretically expected ratios, demonstrating high quantitative accuracy across different gradient lengths.

### Assessing Quantitation in Extreme Ratio Three-Proteome Mixtures

To further challenge the quantitation capabilities of the workflow, we designed an “extreme ratio” three-proteome mixture. In this experiment, digests of *E. coli* and yeast were mixed in varying, inverse proportions against a constant background of 50% of HeLa digest. This simultaneously created samples with both large fold changes for the yeast proteome (from 0.5% to 20%, a 40-fold change) and subtle fold changes for the E. coli proteome (from 49.5% to 30%, a ∼1.6-fold change) (**Figure 2A**). Even with the 300 SPD method, the workflow consistently identified 7,000-8,000 protein groups across all mixtures from a 200 ng total protein load (**Figure 2B**).

**Figure 2.**
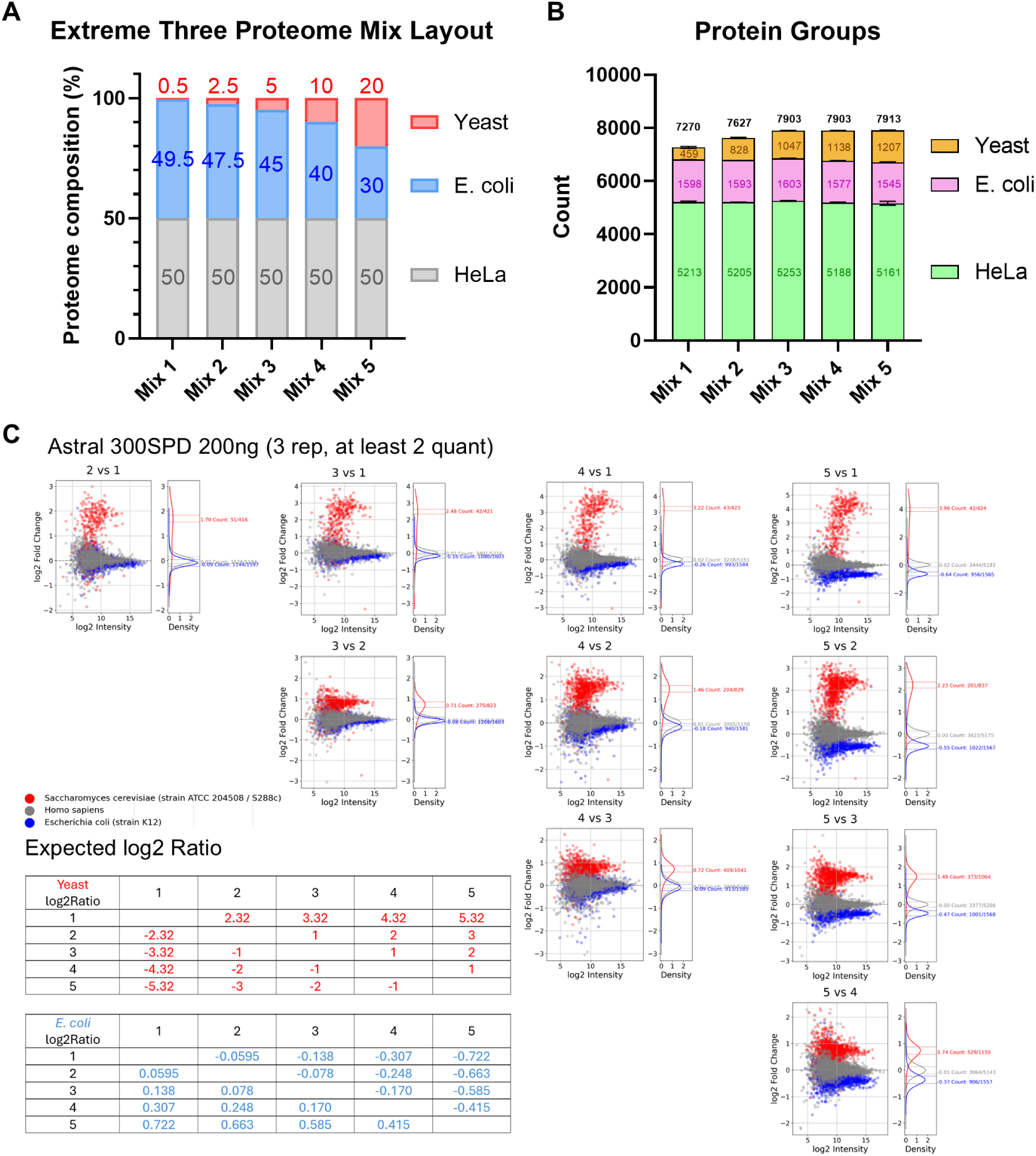
Accurate Quantification of Extreme Ratio Three-Proteome Mixtures. (A) Schematic table detailing the composition of the five extreme-ratio mixtures, with varying amounts of yeast and *E. coli* digest against a constant HeLa background. The expected log_2_Ratio between samples is shown for both yeast and *E. coli*. (B) Bar charts showing the number of protein groups in each mixture sample with a 200 ng loading amount. (C) Scatter plots showing the measured log_2_Ratio versus log_2_(Intensity) for pairwise comparisons between mixtures. Proteins for yeast (red), *E. coli* (blue), and human (gray) are shown. The density plots on the right of each panel illustrate that the maximum density value of the observed fold-change. Data points within 10% error of maximum observed ratio were counted. Three technical replicates were performed.

The workflow showed great linearity in quantifying these mixtures. Remarkably, the subtle fold-changes of the *E. coli* proteome were quantified with high accuracy, closely tracking the expected theoretical ratios (**Figure 2C**, blue dots). As expected, some ratio compression was observed for the large-fold-change yeast proteome, particularly for lower abundance proteins (**Figure 2C**, red dots).The performance was also validated with even lower sample loading amounts (50 ng in **Figure S2A**). Notably, we observed less ratio compression with the longer gradient compared to the results from the shorter gradient experiment shown in Figure 2, particularly for high fold-change comparisons (**Figure S1, Figure S2B**). We also observed less ratio compression when comparing the fold changes on the peptide level (**Figure S3**).

All these observation highlights the ability of LFQ-DIA on Orbitrap Astral mass spectrometer to maintain quantitative accuracy over a wide dynamic range, a critical feature for identifying significant biological changes in complex samples. Crucially, this experiment demonstrates the method’s strength in confidently quantifying small, precise changes even when other proteins in the sample are changing dramatically.

### Development and Validation of a 300 SPD Method Using an 8-cm Column

Building on the initial benchmarks, we further developed an optimized ultra-high-throughput method capable of analyzing 300 SPD (**Table S1**). To achieve this, we utilized a recently released 8-cm IonOpticks Aurora Rapid column, which is compatible with commercial heating devices, ensuring greater chromatographic reproducibility through temperature control. Despite the very short gradient time, this method achieved an impressive 87% usage of the total MS acquisition time, maximizing instrument efficiency (**Figure 3A**).

**Figure 3.**
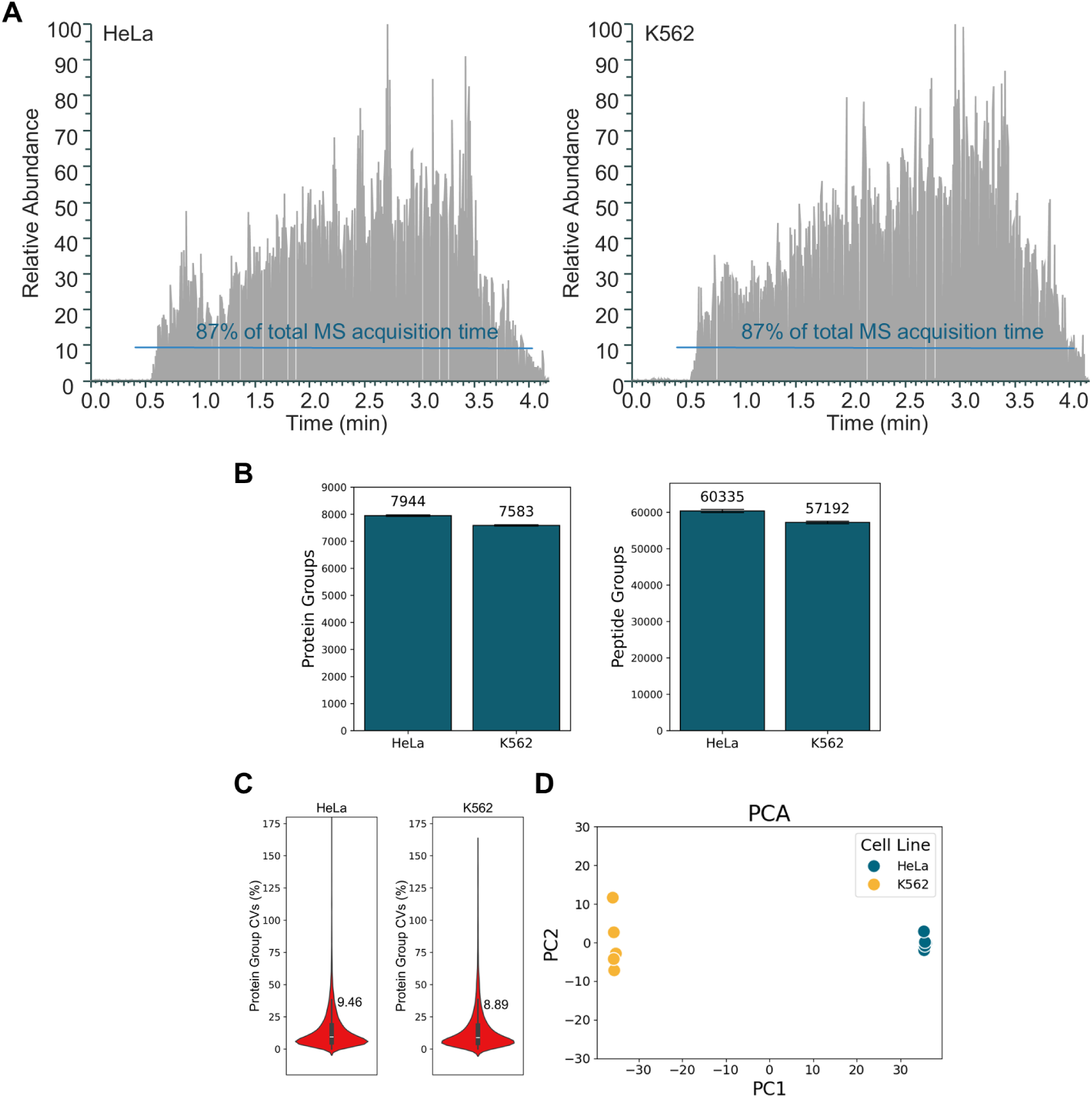
In-depth and Precise Proteome Analysis at 300 SPD. (A) A representative total ion chromatogram from a 200 ng HeLa and K562 digest analyzed with the 4-minute gradient on an 8-cm column. (B) The number of protein and peptide groups identified from HeLa and K562 cells (n=5, mean ± SD). (C) Violin plots showing the distribution of protein group coefficient of variance (CV) percentages from five replicate injections for each cell line, with medians labeled. (D) Principal component analysis (PCA) plot of quantified proteins, showing clear separation between the HeLa and K562 cell lines.

To evaluate the proteome coverage and quantitative precision of this new method, we analyzed 200 ng digests of two different human cell lines, HeLa and K562. The workflow identified 7,944 protein groups in HeLa and 7,583 protein groups in K562, along with over 60,000 and 57,000 peptide groups, respectively, indicating excellent proteome depth for screening applications (**Figure 3B**). Furthermore, the median coefficient of variance (CV) for protein quantification was approximately 9% for both cell lines, demonstrating high precision (**Figure 3C**). This high reproducibility is the foundation for confidently detecting small biological perturbations. A principal component analysis (PCA) showed clear and distinct clustering of the two cell lines, confirming that the method is sensitive enough to distinguish between different biological samples (**Figure 3D**).

### Demonstrating Quantitative Linearity with a Two-Proteome Mixture

After establishing the depth and precision of the 8 cm column 300 SPD method, we assessed its quantitative linearity. For this, we initially performed a two-proteome mixture experiment where increasing amounts of yeast digest (from 20 ng to 160 ng) were spiked into a constant background of 200 ng of HeLa digest (**Figure 4A**). By comparing the measured protein intensities for the yeast proteome across these samples, we observed a strong linear correlation between the signal and the amount of yeast digest spiked in out of the ∼7,000 identified human proteins and ∼3,000 yeast proteins (**Figure S4A, B**). The linear regression analysis yielded an R^2^ value of 0.999, underscoring the outstanding quantitative accuracy and linearity of the 300 SPD method (**Figure 4B**).

**Figure 4.**
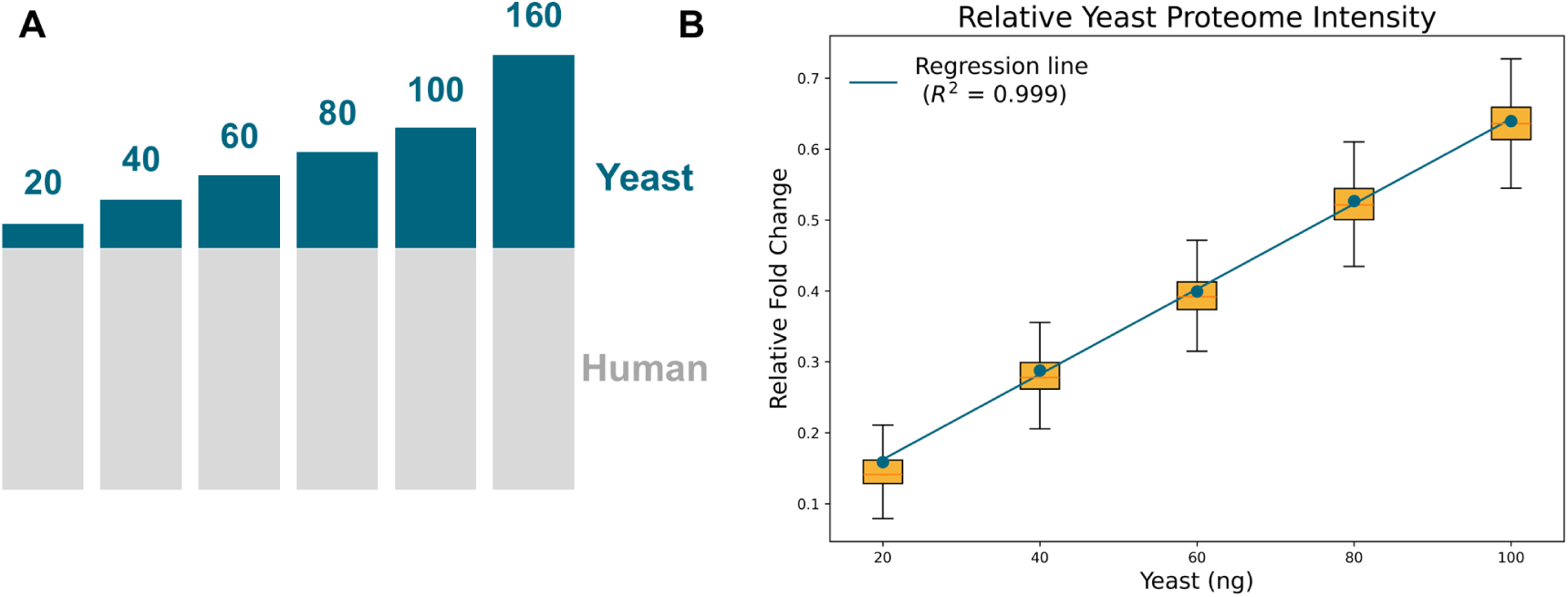
Excellent Quantitative Linearity of the Ultra-High-Throughput Workflow. (A) Schematic diagram of the two-proteome mixture experiment, where 20, 40, 60, 80, 100, and 160 ng of yeast digest were spiked into a 200 ng HeLa digest background. (B) A plot of the relative fold change of the yeast proteome versus the amount of yeast digest added. Boxes show the three quartile values of the distribution. The whiskers extend to points that lie within 1.5 interquartile ranges (IQRs) of the lower and upper quartile. The data points show a clear linear trend, with a linear regression fit demonstrating an R² of 0.999.

### Application in TPD: Profiling Dose-Dependent Effects of CELMoDs

With full confidence in the developed high-throughput method, we applied it to a real-world drug screening scenario: studying the proteome-wide effects of cereblon E3 ligase modulators (CELMoDs) for targeted protein degradation (TPD). We treated human Jurkat T cells for 24 hours with four different concentrations (8, 40, 200, and 1000 nM) of three immunomodulatory drugs (IMiDs): pomalidomide, iberdomide, and mezigdomide (**Figure 5A**).

**Figure 5.**
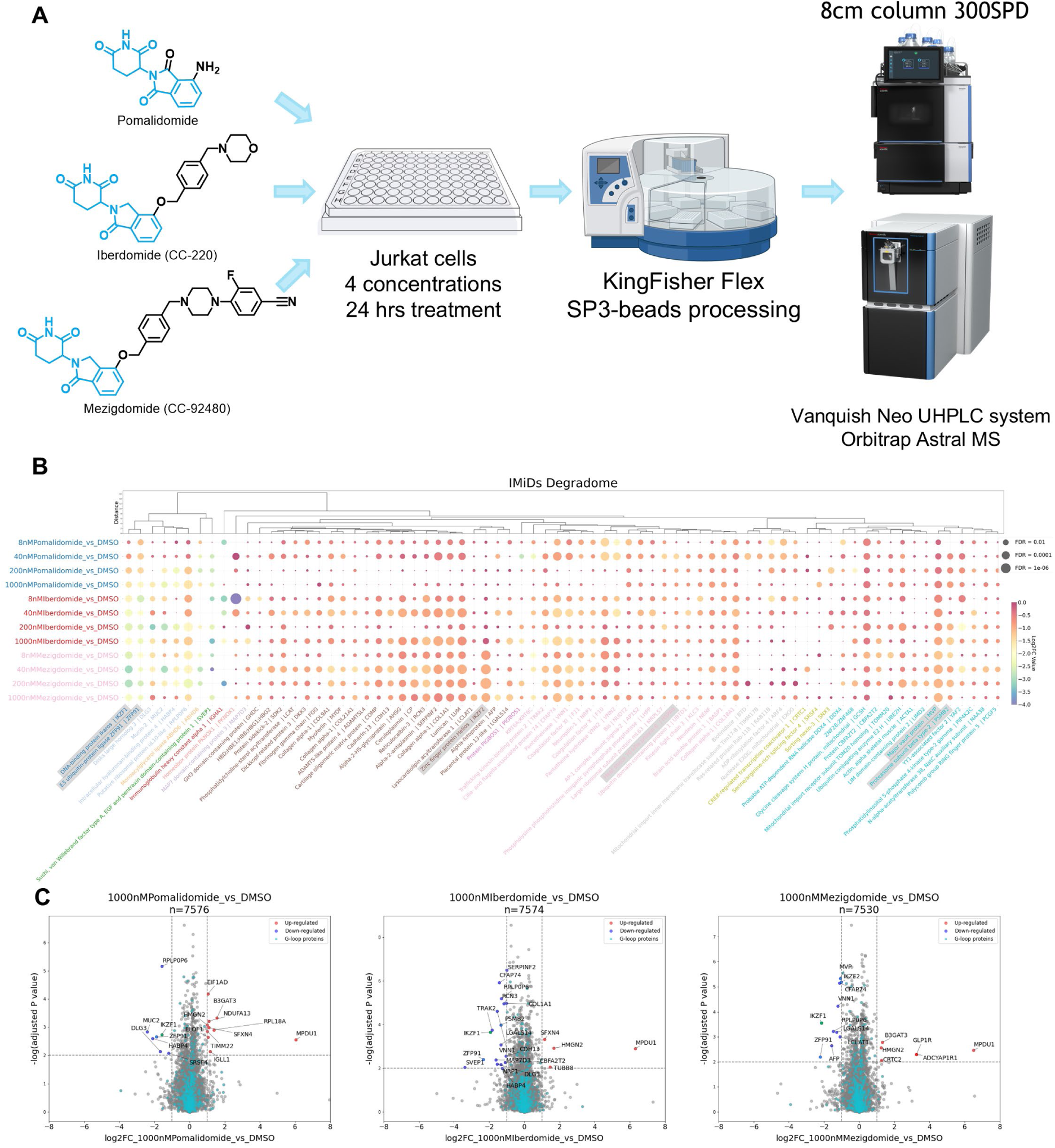
Dose-Dependent Protein Degradation Induced by Pomalidomide, Iberdomide, and Mezigdomide. (A) Schematic diagram showing experimental design. Cells were treated with various concentrations of IMiDs (pomalidomide, iberdomide, and mezigdomide) for 24 hrs, followed by KingFisher-assisted SP3 magnetic beads sample preparation and LC-MS analysis with 8-cm column and 300 SPD method. (B) Bubble plot showing the dose-dependent degradation of IKZF proteins by pomalidomide, iberdomide, and mezigdomide in human Jurkat cells. The bubble color denotes the relative degradation magnitude, while bubble size reflects statistical significance based on adjusted p-values. (C) Volcano plots illustrating the differential protein level analysis in Jurkat cells treated with 1000 nM of pomalidomide, iberdomide, or mezigdomide compared to DMSO-treated controls. (2 biological replicates by 2 technical replicates)

The analysis revealed distinct, dose-dependent protein degradation patterns for each compound (**Figure 5B**). As expected, all three drugs induced the degradation of known cereblon neosubstrates like IKZF1 and ZFP91 (**Figure 5B, C**), with mezigdomide showing the highest potency at lower concentrations. Interestingly, our data revealed that mezigdomide also led to the significant degradation of additional proteins, most notably IKZF2. The degradation of IKZF2 in T cells is thought to enhance anti-cancer immunity ^19^, suggesting that mezigdomide may have enhanced therapeutic efficacy due to its broader substrate profile. These findings demonstrate the power of our ultra-high-throughput workflow for chemoproteomics-based drug screening and discovering novel mechanisms of action. Additional volcano plots for all concentrations are provided in **Figure S5**.

## Discussion

In this study, we successfully developed and validated an ultra-high-throughput proteomics workflow capable of analyzing 300 samples per day using the Orbitrap Astral mass spectrometer. The method delivers deep proteome coverage, identifying nearly 8,000 protein groups from human cell lines with excellent quantitative precision, making it highly suitable for large-scale applications such as compound screening for targeted protein degradation. Our results from the TPD application underscore this, where we not only confirmed known targets of IMiDs but also identified novel dose-dependent degradation patterns, such as for IKZF2 by mezigdomide, providing new biological insights.

A key aspect of validating any quantitative workflow is understanding its limitations, particularly when pushing the boundaries of throughput. Our analysis of the extreme three-proteome mixture was designed specifically to challenge the method’s performance across a wide dynamic range, covering the fold change from ∼2.5% to ∼95%. With the 300 SPD method, we observed some ratio compression, a phenomenon where the measured fold change is smaller than the true biological change. This effect was most pronounced for proteins undergoing large fold changes and was particularly evident for lower-intensity proteins. This is a well-documented challenge in shotgun proteomics especially in isobaric labeling quantification techniques like tandem mass tag (TMT), often attributed to factors like co-isolation of interfering precursor ions, especially in complex samples analyzed with very short chromatographic gradients ^20^. However, for DIA, this issue is substantially mitigated because quantification relies on multiple fragment ions. Modern DIA analysis software can effectively distinguish and quantify target peptides by matching their specific, high-quality fragment ion spectra from the Astral analyzer, even in the presence of co-eluting interferences at the precursor level. Nevertheless, another layer of complexity arises for low-abundance proteins, as their quantification often relies on a small number of identified peptides—sometimes only one or two. This can introduce a degree of stochasticity, making their quantification inherently less robust than that of more abundant proteins represented by multiple peptides.

Significantly, we demonstrated that this ratio compression could be effectively mitigated by adjusting the analytical method. By employing a longer gradient (60 SPD), we observed improvement in quantitative accuracy and a reduction in ratio compression for the same extreme proteome mixture sample. This highlights the flexibility and strength of the Orbitrap Astral platform. The ultra-fast 300 SPD method is ideal for primary screening, where the goal is to rapidly identify the most potent hits and significant changes. For subsequent validation or more detailed follow-up studies requiring higher quantitative fidelity for low-abundance targets, switching to a moderately longer gradient provides a straightforward solution without compromising the core benefits of the platform.

Furthermore, the exceptional sensitivity of the Astral analyzer is particularly powerful for resolving small fold changes. This capability is invaluable in early-stage drug discovery campaigns, where initial compound hits may only induce small, yet statistically significant, changes in protein abundance. The robustness of this approach was validated under stringent, blind-test conditions. The three-proteome mixture experiment was conducted as a blinded evaluation where the sample ratios, prepared by BCM, were unknown to Thermo Fisher scientists during data acquisition and analysis. The subsequent accurate determination of the proteome ratios against the ground truth demonstrated the platform’s reliability and convinced us that it meets the highly demanding requirements for drug discovery workflows in chemoproteomics ^21–23^ and targeted protein degradation ^1,2,4,24,25^.

In conclusion, the Orbitrap Astral mass spectrometer enables a paradigm shift in high-throughput proteomics. The workflow presented here offers a powerful, scalable, and adaptable solution for drug discovery and other large-scale proteomics studies. While ratio compression of low-intensity proteins is a consideration in ultra-fast methods, we have shown that it is a manageable variable, allowing researchers to strike the optimal balance between throughput and quantitative depth to meet diverse experimental needs.

## Acknowledgement

The research was supported in part by the Michael E. DeBakey, M.D., Professorship in Pharmacology and the seed funding for Center for NextGen Therapeutics.

## Supplementary Figures

**Figure S1.**
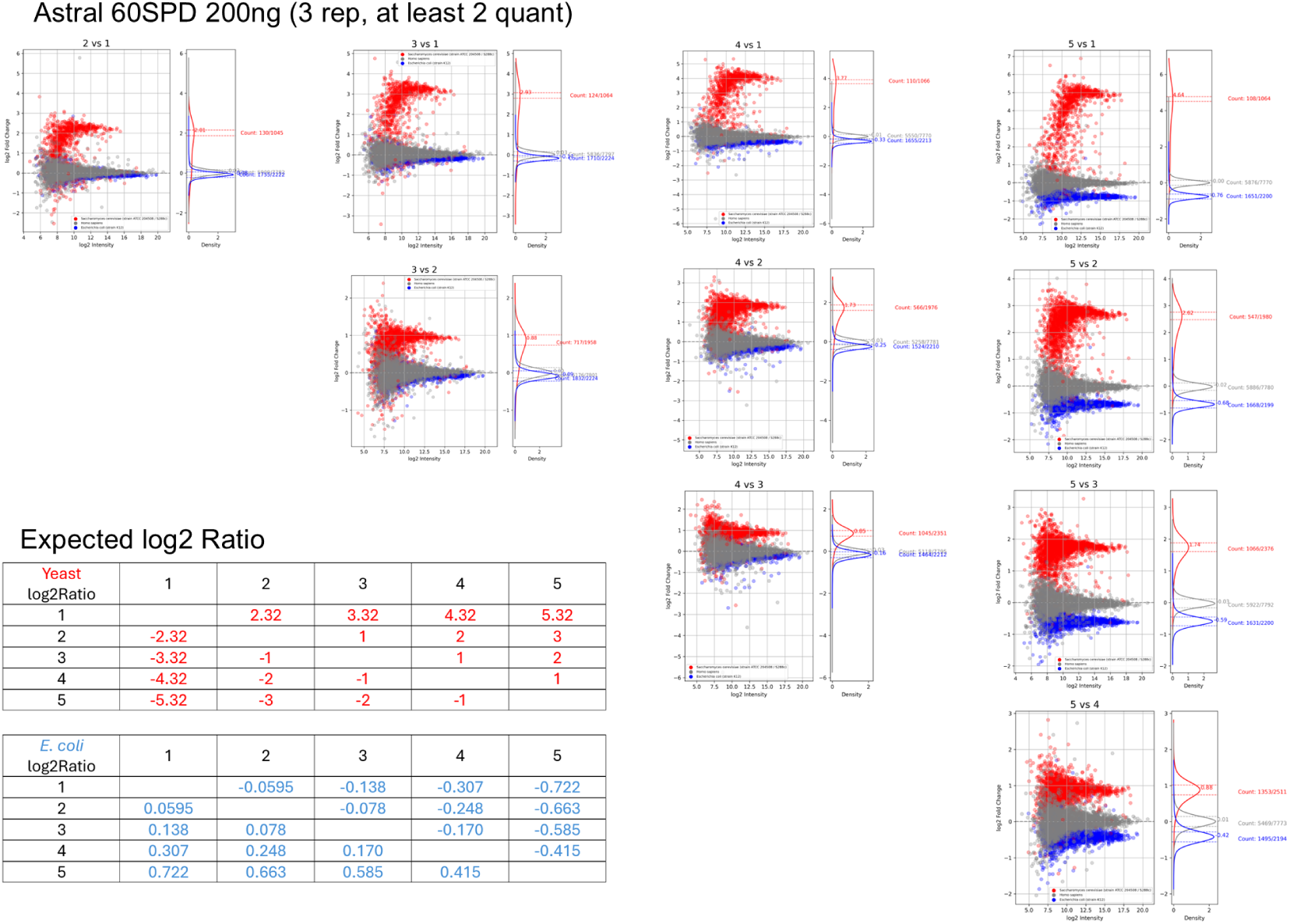
With 200 ng loading amount of extreme three-proteome mix and 60 SPD method, individual protein ratio was plotted against protein intensity. Ratio distribution was calculated with Gaussian-KDE and data points within 10% error of maximum observed ratio were counted. Three technical replicates were performed.

**Figure S2.**
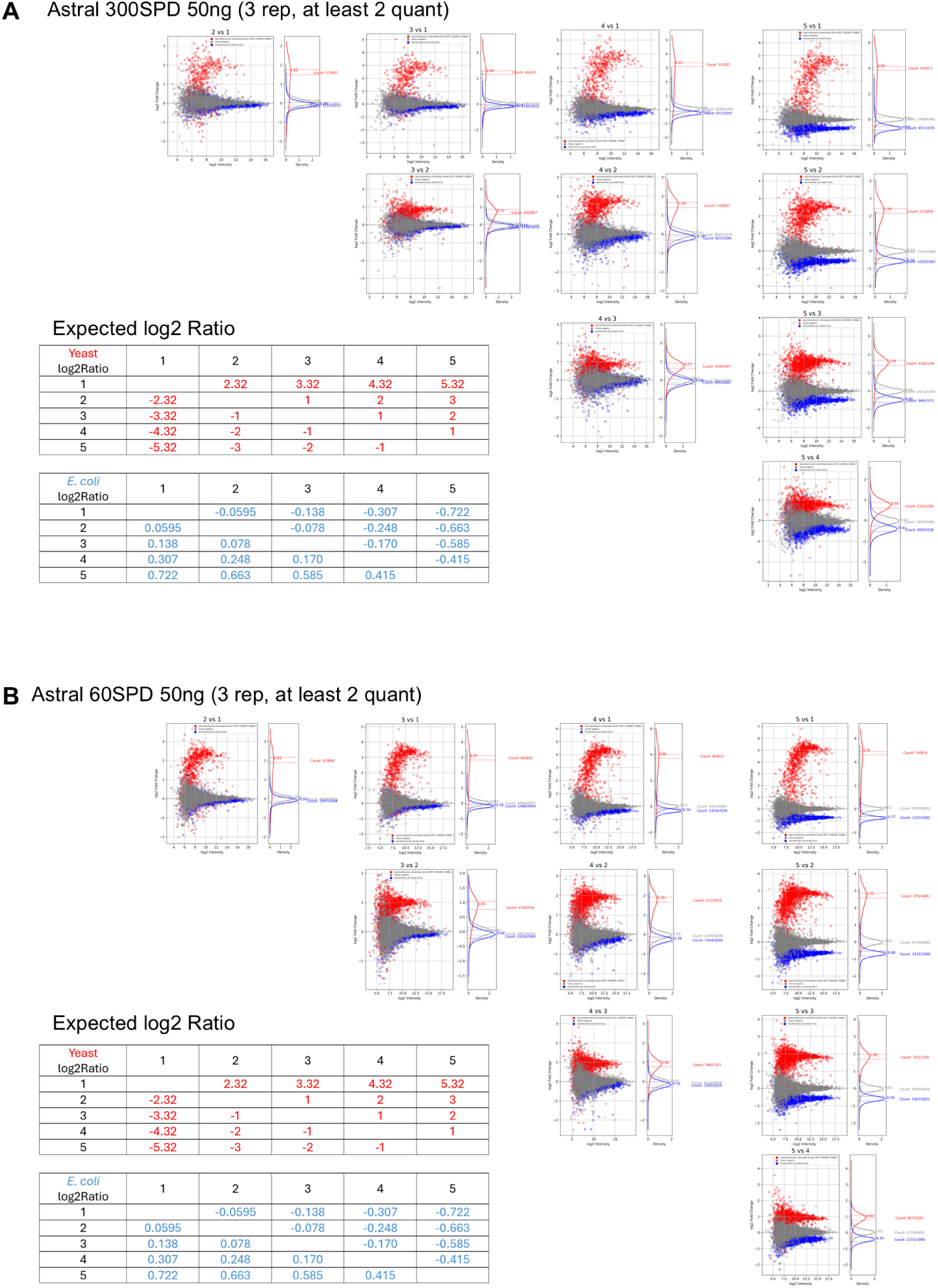
Accurate quantification of LFQ-DIA workflow on low loading sample (50 ng). With 50 ng loading amount of extreme three-proteome mix and 300 SPD (5 cm column) (A) or 60 SPD (8 cm column) (B) method, individual protein ratio was plotted against protein intensity. Ratio distribution was calculated with Gaussian-KDE and data points within 10% error of maximum observed ratio was counted. Three technical replicates were performed.

**Figure S3.**
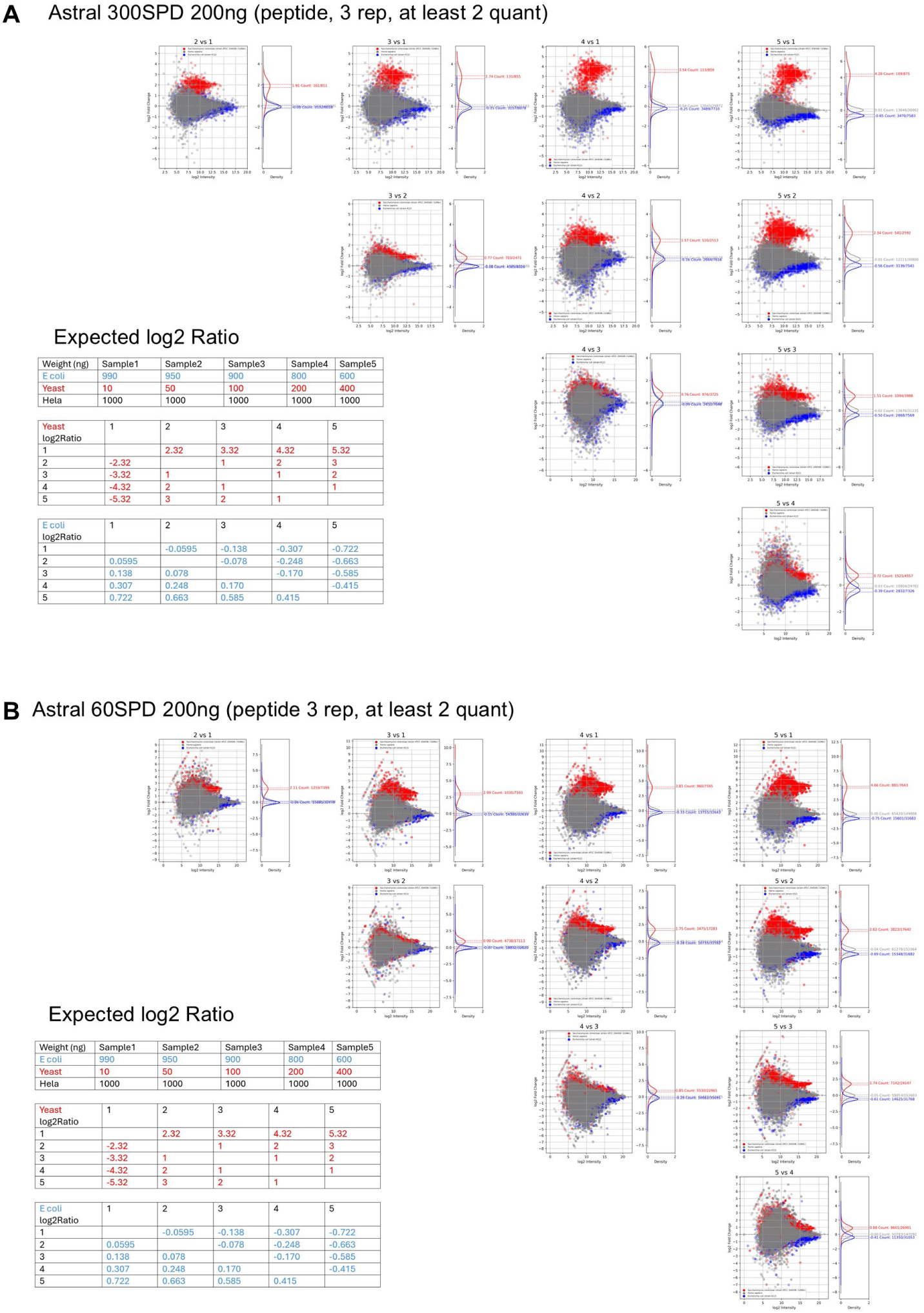
Peptide group quantification of LFQ-DIA workflow on high loading sample (200 ng). With 200 ng loading amount of extreme three-proteome mix and 300 SPD (**A**) or 60 SPD (**B**) method, individual peptide ratio was plotted against peptide intensity. Ratio distribution was calculated with Gaussian-KDE and data points within 10% error of maximum observed ratio were counted. Three technical replicates were performed.

**Figure S4.**
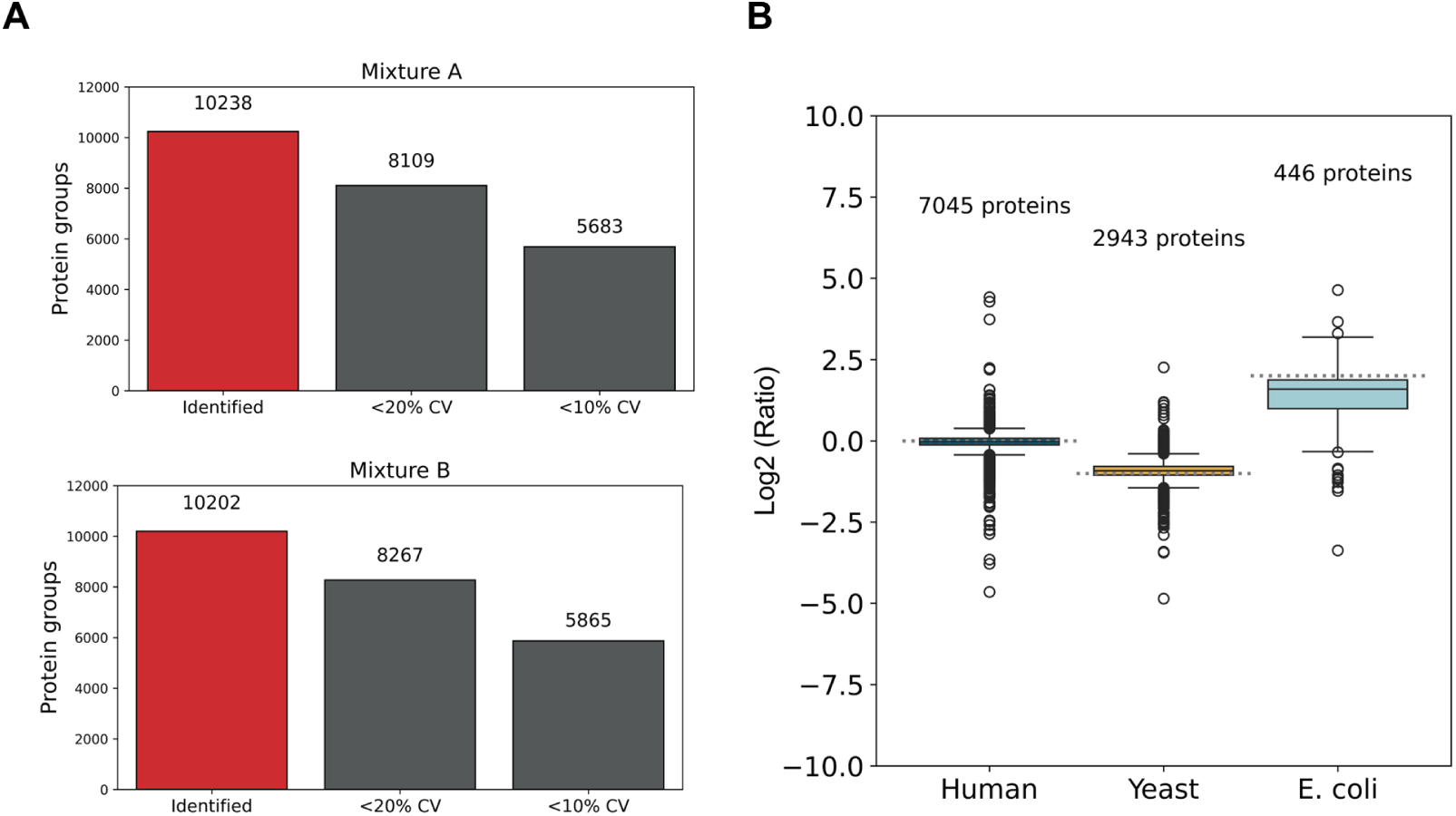
Accurate quantification linearity of the 8 cm column LFQ-DIA workflow. Standard three-proteome mix (Figure 1B) was loaded onto 8-cm column and analyzed with the 300 SPD method. (A) Identified protein group numbers with different CV of quantitation cutoff. (B) Quantification and quantitation accuracy test using standard three proteome mixtures. Boxes show the three quartile values of the distribution. The whiskers extend to points that lie within 1.5 IQRs of the lower and upper quartile. Dash lines indicate the expected ratio values for each proteome.

**Figure S5.**
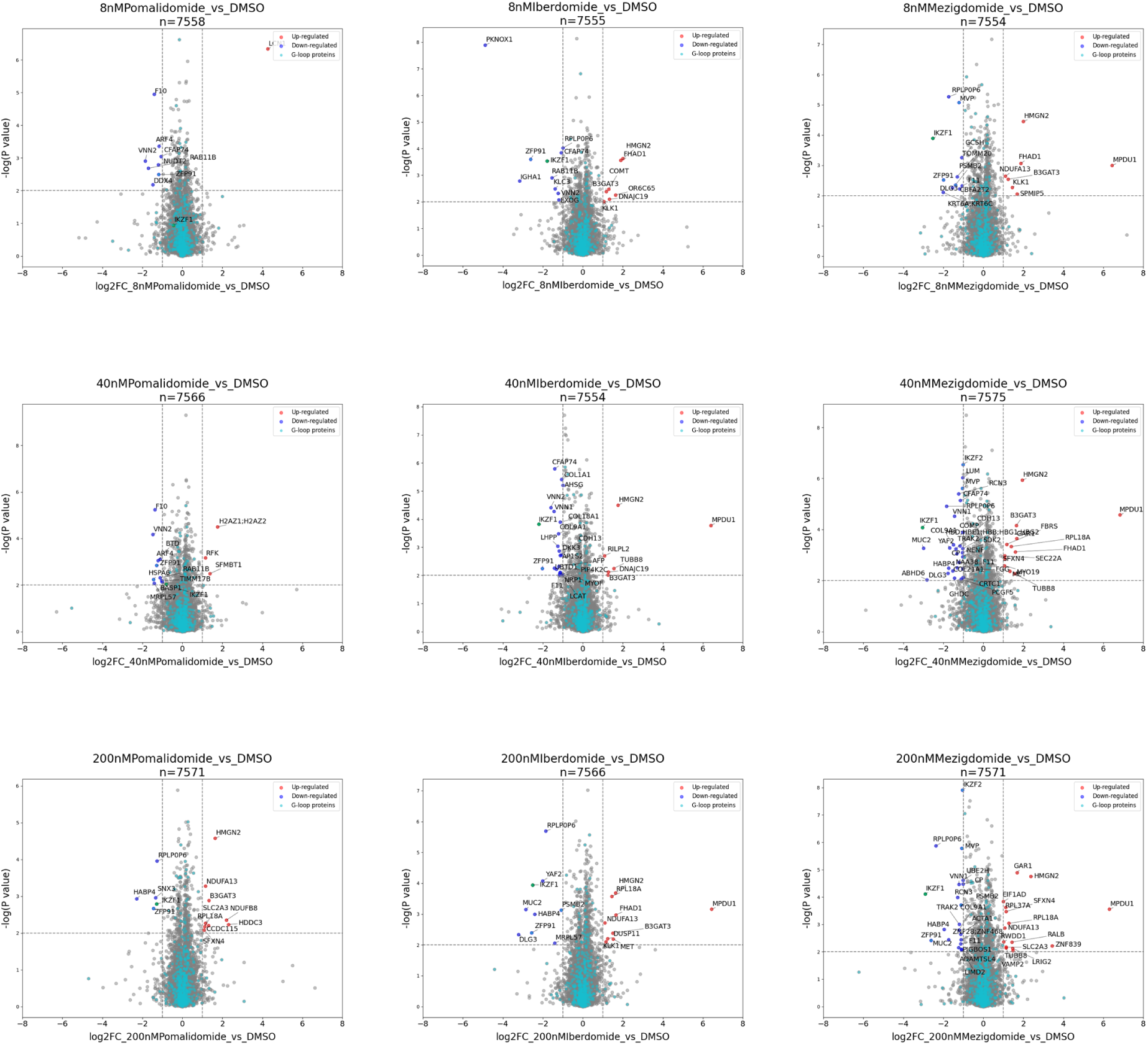
Volcano plots illustrating the differential protein level analysis in Jurkat cells. Jurkat cells were treated with 8, 40, 200 nM of pomalidomide, iberdomide, or mezigdomide and compared with DMSO-treated controls. (2 biological replicates by 2 technical replicates)

